# A morphological trait involved in reproductive isolation between Drosophila sister species is sensitive to temperature

**DOI:** 10.1101/2020.01.20.911826

**Authors:** Alex E. Peluffo, Mehdi Hamdani, Alejandra Vargas-Valderrama, Jean R. David, François Mallard, François Graner, Virginie Courtier-Orgogozo

**Affiliations:** Institut Jacques Monod, CNRS, Univ. de Paris, 75013 Paris, France; Institut Systématique Evolution Biodiversité (ISYEB), CNRS, MNHN, Sorbonne Université, EPHE, 57 rue Cuvier, CP 50, 75005 Paris, France; Laboratoire Evolution, Génomes, Comportement, Biodiversité (EGCE), CNRS, IRD, Univ. Paris-sud, Université Paris-Saclay, 91198 Gif-sur-Yvette, France; Institut de Biologie de l’École Normale Supérieure, CNRS UMR 8197, Inserm U1024, PSL Research University, F-75005 Paris, France; Matière et Systèmes Complexes, CNRS UMR 7057, Univ. de Paris, Paris, France

**Keywords:** Plasticity, Speciation, Genitalia, Reproductive isolation, Drosophila, Landmark, Automatic Detection, Shape analysis

## Abstract

Male genitalia are usually extremely divergent between closely related species, but relatively constant within one species. Here we examine the effect of temperature on the shape of the ventral branches, a male genital structure involved in reproductive isolation, in the sister species *Drosophila santomea* and *D. yakuba*. We designed a semi-automatic measurement pipeline that can reliably identify curvatures and landmarks based on manually digitized contours of the ventral branches. With this method, we observed that temperature does not affect ventral branches in *D. yakuba* but that in *D. santomea* ventral branches tend to morph into a D. yakuba-like shape at lower temperature. Our results suggest that speciation of *D. santomea* and *D. yakuba* was associated with a change in genitalia plasticity.

## Introduction

Phenotypic plasticity, the capacity for one genotype to generate multiple phenotypes in response to environmental variation, is a pervasive feature of biological systems (Debat & David, 2001; Klingenberg, 2019). The connection between plasticity and speciation is multifaceted (Lafuente & Beldade, 2019). On the one hand, plasticity can be heritable and modified by selection. On the other hand, plasticity can favor adaptation and speciation. As animals colonize novel habitats or face changing climate conditions, the phenotypic traits that are optimal for fitness are usually different from those experienced in the ancestral population. Waddington was among the first to suggest that organisms may solve this challenge by phenotypic plasticity first and later on by genetic fixation of what was previously an environmentally induced phenotypic trait (a process he called "genetic assimilation") (Waddington, 1942). According to several authors, the trait variations enabled by plasticity can initiate and accelerate the pace of adaptive evolution and promote morphological diversification. This central idea is at the basis of the “flexible stem hypothesis” (West-Eberhard, 2003; Schneider & Meyer, 2017) the “plasticity-first” model (Levis & Pfennig, 2016), where plasticity allows the population to persist long enough for adaptive mutations to arise and become fixed. A key feature of all these views is that the phenotypic change triggered by the plastic response, which allows the colonization of the new niches, is a phenocopy, i.e., that the phenotypic change can be developmentally triggered by environmental variation or genetic variation interchangeably (Lafuente & Beldade, 2019). As we learn more about the genes mediating phenotypic plasticity (Gibert, 2017), it appears that similar phenotypic changes, either environmentally or genetically induced, can sometimes involve the same genetic loci. For example, the same enhancer of the gene *tan* contributes to both phenotypic plasticity in *Drosophila melanogaster* (Gibert *et al.*, 2016) and interspecific evolution between sister-species *D. santomea* and *D. yakuba* with respect to abdomen pigmentation (Jeong *et al.*, 2008).

Depending on the setting, plasticity can either accelerate, slow down, or have little effect on evolution and species divergence (Price *et al.*, 2003). Speciation, the process through which lineages diverge and become reproductively isolated, involves the accumulation over time of barriers limiting interbreeding, including divergence in ecological niches, behavioral isolation and genomic incompatibilities (Coyne & Orr, 2004). As early as 1844, anatomical differences in genitalia between closely related species were proposed to be an essential mechanism maintaining reproductive isolation, as the so-called “lock-and-key” hypothesis (Dufour, 1844; Masly, 2011). In animals with internal fertilization, genitalia are the most rapidly evolving organs in terms of morphology (Eberhard, 1988), suggesting that a significant part of the speciation process involves anatomical divergence in genitalia. Alternatively, genital evolution can be a by-product of other evolutionary processes occurring within single lineages, independently of speciation (such as sexual selection), and lead to reproductive isolation as a by-product, when individuals attempt to hybridize with other lineages (Masly, 2011).

The lock-and-key hypothesis, even in species where it seems applicable, has been challenged by a variety of observations, including the facts that (1) genitalia in females do not differ as much as in males, (2) closely related species with conspicuous genital differences can still often produce hybrids, (3) males with laser-ablated genital organs can still copulate with no observed defect, and (4) genitalia morphology can be sensitive to temperature or nutrition (Shapiro & Porter, 1989; Arnqvist & Thornhill, 1998; Andrade *et al.*, 2005; Masly, 2011; Simmons, 2014; LeVasseur-Viens *et al.*, 2015)and references therein). It is thus possible that in some taxonomic groups interspecific differences in genital morphology do not contribute much to reproductive isolation.

To better comprehend the link between plasticity and speciation, careful examinations of particular cases are essential, and genital traits involved in reproductive isolation represent highly relevant model systems. How plastic are genitalia in general? Surprisingly, few studies have examined genitalia after raising organisms in various conditions. In the water strider *Aquarius remigis*, the mosquito *Aedes aegypti* and the fly *D. melanogaster*, changes in larval crowding, nutrition conditions or temperature were found to affect adult body size but had little effect on the size of the external genitalia (Wheeler *et al.*, 1993; Fairbairn, 2005; Shingleton *et al.*, 2009). However, in two other species, the mosquito *Anopheles albimanus* and the fly *Drosophila mediopunctata*, the size and shape of the male intromittent organ was found to vary with rearing temperature (Hribar, 1996; Andrade *et al.*, 2005). Overall, analysis of individuals sampled from the wild show that for a given arthropod or mammal species, the genitalia are usually more or less the same size whereas adult body size varies extensively (Eberhard *et al.*, 1998; Dreyer & Shingleton, 2011 and references therein). These observations are concordant with the “lock-and-key hypothesis”, where male genitalia have to be of a particular size and shape to physically fit with the female genitalia. They are also explained by the “one-size-fits-all” hypothesis, where females appear to prefer males with genitalia of intermediate size (Eberhard *et al.*, 1998).

In order to analyze and quantify the possible link between plasticity, reproductive isolation and interspecific divergence, we chose to examine the effect of temperature on a male primary sexual trait likely involved in reproductive isolation between two Drosophila sister species, *D. santomea* and *D. yakuba*. These two species form an attractive system because their natural environment is relatively well characterized, they are known to hybridize, and one of their most remarkable morphological differences is a primary sexual trait that seems to be involved in a “lock-and-key” mechanism. *D. santomea* and *D. yakuba* diverged approximately 0.5-1 million years ago (Turissini & Matute, 2017). They can be crossed to generate fertile F1 females (Lachaise *et al.*, 2000). *D. santomea* is endemic to the island of São Tomé, a volcanic island off the coast of Gabon (Lachaise *et al.*, 2000), while *D. yakuba* is found in São Tomé and throughout sub-Saharan Africa (Lachaise *et al.*, 1988, 2000). In São Tomé, *D. santomea* lives in the mist forests at high elevations while *D. yakuba* is found in open habitats associated with human presence, mostly at low elevations (Llopart *et al.*, 2005a; b) Both species co-occur at mid-elevation, around 1150 m, and hybrids have been found consistently in this hybrid zone since its discovery in 1999 (Lachaise *et al.*, 2000; Llopart *et al.*, 2005a; Comeault *et al.*, 2016; Cooper *et al.*, 2018). *D. santomea* being insular, it is thought that this species originated from a common ancestor with *D. yakuba*, which colonized the island about 0.5-1 million years ago (Cariou *et al.*, 2001; Llopart *et al.*, 2002; Turissini & Matute, 2017), and that the present co-occurence of *D. santomea* and *D. yakuba* in São Tomé reflects secondary colonization by *D. yakuba* from the African mainland, maybe during the last 500 years when Portuguese colonised the island (Cariou *et al.*, 2001). Analysis of genomic and mitochondrial DNA sequences indicate that gene flow occurred between the *D. santomea* and *D. yakuba* more than 1,000 generations ago (Turissini & Matute, 2017; Cooper *et al.*, 2019).

Multiple potential reproductive isolating mechanisms have been identified between the two species, such as genetic incompatibilities (Coyne *et al.*, 2004; Moehring *et al.*, 2006), ecological niche divergence (Matute *et al.*, 2009), mate discrimination (Lachaise *et al.*, 2000; Coyne *et al.*, 2002), behavioral (Cande *et al.*, 2012), physiological (Matute, 2010) and morphological differences (Lachaise *et al.*, 2000; Jeong *et al.*, 2008; Nagy *et al.*, 2018; Liu *et al.*, 2019). One reproductive isolating mechanism between *D. yakuba* and *D. santomea* involves a difference in ventral branches shape in the male genitalia and is the most conspicuous difference in male genitalia shape between the two species (Kamimura & Mitsumoto, 2012b; Yassin & Orgogozo, 2013) (Figure 1). Ventral branches are located below the aedeagus [i.e., the insect phallus (Rice *et al.*, 2019)] and are only found in the *D. yakuba* complex, which comprises *D. teissieri*, *D. yakuba* and *D. santomea* (Yassin and Orgogozo 2013).

**Figure 1.**
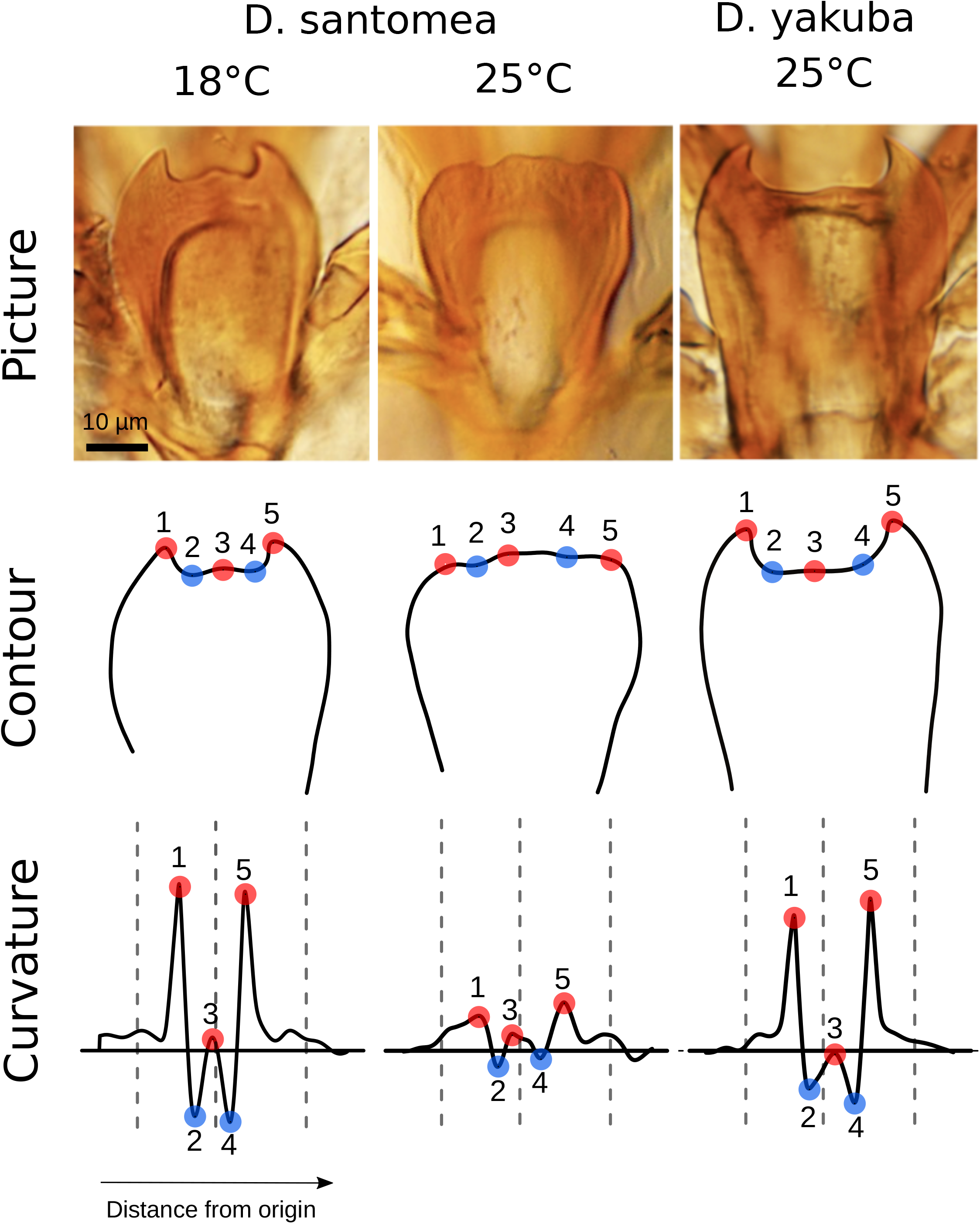
Landmark detection for ventral branches at 18°C and 25°C. For each individual, a picture of the ventral branches is taken (top panel). The contour is digitized by hand and smoothed (middle panel). The curvature along the contour is obtained by finite differences, which are iterated for refining, and smoothed too (bottom panel). Smoothed curvature (vertical axis), measured in inverse micrometers, is plotted along the contour, starting from the leftmost point. The horizontal axis is the distance along the contour, called the curvilinear abscissa, measured in micrometers. Here both axes are normalized by size and represented in arbitrary units. In each plot the left dashed vertical line is the automatic detection lower bound, the middle dashed line is the imputed global midpoint, the right dashed line is the automatic detection upper bound (see methods). Red points represent peaks and therefore curvature maxima whereas blue points represent cavities and therefore curvature minima. Since *D. yakuba* is not sensitive to temperature, we only show one characteristic shape. To understand where this genital structure is positioned within the male genitalia, see Fig. 1 of Kamimura & Mitsumoto, 2012b.

In *D. yakuba*, spiny ventral branches insert inside female protective pouches during mating. In *D. santomea*, the male spines and female pouches are absent. These structures appear to play important roles during copulation. When mating with *D. yakuba* males, *D. santomea* females are wounded by the spines of the male ventral branches and they live shorter than females mating with conspecific males (Matute & Coyne, 2010; Kamimura, 2012; Kamimura & Mitsumoto, 2012b). Compared to *D. teissieiri* females, *D. santomea* females also survive less to interspecific copulation with *D. mauritiana* (Yassin & David, 2016). Moreover, Kamimura and Mitsumoto (2012b) reported that “copulating pairs of *D. santomea* males x *D. yakuba* females dislodge readily when disturbed”, suggesting that the spines may fasten genital coupling (Masly, 2011). We previously found that a major QTL on chromosome 3L contributes to the ventral branches shape difference between *D. santomea* and *D. yakuba* (Peluffo *et al.*, 2015).

In São Tomé the climate is very stable throughout the year, with only a 2.5°C-difference between the average daily temperature of the warmest month (March) and of the coldest one (July), and daily oscillations of about 5°C only (climate-data.org, www.worldclim.org/bioclim). Based on temperature measurements at Monte Café (climate-data.org), we estimate that the average temperature in the hybrid zone of Bom Sucesso (1153m) varies between 15.5°C and 18°C throughout the year. In the wild, *D. santomea* flies are thus likely developing mainly at temperatures around 18°C or lower.

In previous studies of ventral branch shape, flies were raised either at 21°C (Yassin & Orgogozo, 2013) or 25°C (Kamimura, 2012; Kamimura & Mitsumoto, 2012b; Peluffo *et al.*, 2015). Here, we report that *D. santomea* males raised at 18°C develop spiny ventral branches comparable to those of *D. yakuba* raised at 25°C. This is a surprising example where organs potentially directly linked with reproductive isolation undergo a plastic modification similar to the difference between two sister species. To better characterize the morphological change in ventral branches shape, we developed a user-friendly method to quantify contour curvatures and automatically detect spines using machine learning. We used it to examine the plastic response of ventral branches development at 18°C and 25°C both in newly collected wild strains and in strains kept in the lab for many years.

## Material and Methods

### Fly rearing and imaging

Fly strains (Table 1) were kept at 22°C on standard yeast-cornmeal–agar medium in uncrowded conditions before the beginning of the experiments. For each strain, roughly 20 individuals were transferred from the 22°C stock to either 18°C or 25°C, kept for a minimum of two non-overlapping adult generations. Adult males were 5 to 7 days old when frozen at −80°C for subsequent dissection. Dissection of genetalia was performed in 1X PBS at room temperature. Each genetalia was mounted as standard glass slides in DMHF (Dimethyl Hydantoin Formaldehyde, Entomopraxis) medium and kept overnight before imaging on an Olympus IX83 inverted station at 40X.

**Table 1.** List of Isofemale lines used in this study. For each species, the most common name in the literature, the location, year of capture and reference to origin of the strain are given. All lines are indicated in the same order as in Figure 2.

| Species | Name | Location | Year | Reference |
| --- | --- | --- | --- | --- |
| *D. yakuba* | Ivory Coast | Ivory Coast | 1955 | Cornell National Drosophila Species Stock Center, Strain #14021-0261.00 (given by D. Stern) |
| *D. yakuba* | BM2015 | São Tomé,<br>Bom Sucesso Botanical Garden,<br>1150m | February 2015 | This study |
| *D. yakuba* | Oku | Cameroun, Mt. Oku, 2000m | April 2016 | This study |
| *D. yakuba* | Raphia | Cameroun, Mt. Oku,<br>1800m | April 2016 | This study |
| *D. santomea* | STO.4 | São Tomé,<br>Obo Natural Reserve, submontane forest<br>1300-1450m | 1998 | (Lachaise *et al.*, 2000)<br>Cornell National Drosophila Species Stock Center, Strain #14021-0271.00 (given by D. Stern) |
| *D. santomea* | STO Cago 1482 | São Tomé<br>1482m | 2001 | (Llopart *et al.*, 2005b)<br>This strain's original name is STO-LAGO 1482 (given by D. Stern) |
| *D. santomea* | Quija22 | São Tomé,<br>Quija River,<br>650m | 2009 | (Gavin-Smyth & Matute, 2013)<br>This strain name is also Quija650.22 (given by D. Matute) |
| *D. santomea* | BM152 | São Tomé<br>Bom Sucesso Botanical Garden,<br>1150m | February 2015 | This study |
| *D. santomea* | BM153 | São Tomé<br>Bom Sucesso Botanical Garden,<br>1150m | February 2015 | This study |
| *D. santomea* | BM161 | São Tomé<br>Bom Sucesso Botanical Garden,<br>1150m | September 2016 | This study |
| *D. santomea* | BM167 | São Tomé<br>Bom Sucesso Botanical Garden,<br>1150m | September 2016 | This study |
| *D. santomea* | 1563 | EYFP lab strain derived from<br>STO CAGO 1482,<br>Insertion @ 3L:11.843.137 | 2001 | (Stern *et al.*, 2017)<br>(given by D. Stern) |

### Raw contour acquisition

All contours were digitized by the same person. Pictures were anonymized for manual contour acquisition so that the digitizer did not know the genotype. Digitization was skipped when the quality of the mounting was judged to be poor. For each picture, a custom ImageJ plugin was used to extract *x*,*y* coordinates (in pixels) of the contour. The plugin is designed to open all the pictures contained in a directory, allowing the user to manually draw a contour of the object of interest using the freehand tool of ImageJ.The raw contour is a series of points *p*_*1*_, *p*_*2*_, …, *p*_*n*_ in a two-dimensional space *x,y* where *n* is the number of points over which the contour passes (usually 500 < *n* < 1000). The contour is open and its endpoints are unimportant (Figure 1). It is analyzed (and twice smoothed) as follows.

### Smoothed contour

The first layer of transformation is a rectangular smoothing filter over the raw contour to obtain the smoothed contour. At each point *p*_*j*_ with coordinates (*x*_*j*_, *y*_*j*_), we derive *p*′_*j*_ with coordinates (*x*′_*j*_, *y*′_*j*_) where 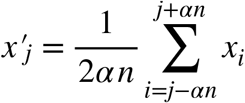and 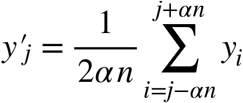.

Here the contour smoothing parameter *α*, to be adjusted via learning, describes the proportion of points (relative to the total number of points forming the contour) to include in the smoothing. This implies that the smoothed contour is *2αn* points shorter (*αn* on each side) than the raw contour.

### Raw curvature of the smoothed contour

For each smoothed contour, the raw curvature *k* is computed with a sliding window of three points. For any set of three points *M*, *N*, *P* forming a triangle, the diameter of the circumscribed circle to this triangle, 2*r* = *MP* / sin (*MN*,*NP*) can be computed as the product of the Euclidean distances divided by the cross product of the two (non-basal) sides, *r* is the curvature radius in *N*, and the curvature *k* in *N* is the inverse of *r* :

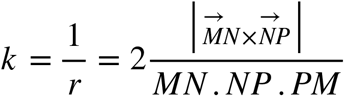

The flatter the contour, the wider the circumscribed circle, the larger the radius *r*, and the smaller the curvature *k*. For each contour, the curvature profile is the curvature *k*_*j*_ computed over *p’*_*j*_ in *p*_′1_, *p*_′*n*_ using its neighbouring points (*M*, *N*, *P* =*p’*_*j*−1_, *p’*_*j*_, *p’*_*j* +1_) versus the curvilinear abscissa *s*_*j*_ of *p’*_*j*_ which is the sum of Euclidean distances from origin, *s*_*j*_ = *p*’_*1*_*p*’_*2*_ + *p*’_*2*_*p*’_*3*_ + … + *p*’_*j*−*1*_*p*’_*j*_.

### Refined curvature

We then use this first raw curvature estimation as information to refine the curvature in a second pass. In this second measure, the refined curvature *k*_*j*_’ is computed over an adaptive window of size 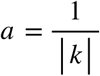 for *k* < 0.1 and *a* = 10 otherwise: *M*, *N*, *P* = *p’*_*j-a*_, *p’*_*j*_, *p’*_*j*+*a*_. This means that the curvature is computed over a larger distance where it is small (and curvature radius is large), which requires more smoothing, without losing the sharpness of curvature peak determination where the curvature is large.

### Smoothed curvature

To improve curvature signal to noise ratio, for each point *p*′_*j*_ with coordinates (*x*′_*j*_, *y*′_*j*_) and refined curvature *k*′_*j*_, we compute the smoothed curvature *k”_j_* as a weighted moving average with triangular weights:

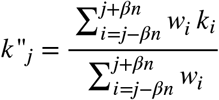

with *w*_*j*_ = *βn*, … , *w*_*i*_ = *βn* − *i* − *j* ∨, … , *w*_*j*−*βn*_ = *w*_*j*+*βn*_ = 0 and where *β* is the smoothing parameter to adjust via learning. *β*describes the proportion of points (relative to the total number of contour points) to include in the smoothing. This implies that the smoothed curvature contour is 2*βn* points shorter (*βn* on each side) than the smoothed contour.

### Landmark detection

Curvature around the start and end of the contour is noisy; it corresponds to a region of low curvature, at the beginning and end of the contour, outside of the region where we expect to find the five landmarks (Figure 1). In addition, the contour digitization by the user, which tends to start at a precise point and to end in a long stroke, results in a slight left-right asymmetry in the curvature profile. After superimposing all smoothed curvature profiles, we choose to exclude the first and last 20% of the smoothed contour. We find that the axis of symmetry (midline) is at position 0.475 instead of 0.5 for a symmetric profile.

Landmarks are Bookstein’s type 2 (local maxima of curvature) (Bookstein, 1992): maxima of the smoothed curvature for landmarks 1, 3, 5 and minima for landmarks 2 and 4. Having detected all minima and maxima, we first define landmark 3 as the maximum closest to the midline position, landmark 2 as the lowest minimum to the left of landmark 3 and landmark 1 as the maximum closest to landmark 2. Following the same logic, we define landmark 4 as the lowest minimum to the right of landmark 3 and landmark 5 as the maximum closest to landmark 4. Having detected all five landmarks, we found that there can seldom be more than one maximum between landmark 2 and landmark 4. In such a situation, we allow resampling of landmark 3 to the highest maximum between landmarks 2 and 4. Finally, we exclude individuals that do not display all five landmarks after detection.

### Spine thrust measure

Having detected all five landmarks, we quantify form using a measure previously introduced (Peluffo *et al.*, 2015), which is highly correlated to the Procrustes analysis principal component measure of inter-specific form variation, and which we called “spine thrust” (ST). ST is a measure of how much spines are elevated above the central ridge of the ventral branches and is computed as:

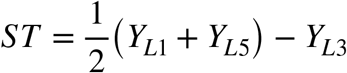

where *Y*_*L*1_, *Y*_*L*3_ and *Y*_*L5*_ are the *Y* coordinate of landmarks 1, 3, 5, respectively. This measurement depends on the precise definition of *X* and *Y* axes. Here the *X*-axis is defined as the axis passing by landmarks 2 and 4 and oriented from 2 to 4, and with the *Y*-axis defined so that (*X*, *Y*) is an oriented orthonormal basis.

### Machine learning

Detection of maxima and minima is a simple feature detection that relies on the derivative of the smoothed curvature profile. However, there are two parameters *α*, *β*, one for each smoothing filter (contour and curvature), which modulate the number and position of these detected maxima and minima. It is possible to explore a set of values for *α* and *β* such that the correlation between manually digitized landmarks and automatically detected landmarks is optimized. Given that humans may introduce bias in the positioning of the landmarks (e.g. if one unconsciously amplifies spine thrust in *D. yakuba* relative to *D. santomea*), the human output may not be optimal over the machine output. This is why we chose not to quantify the learning success rate of our algorithm using the Area Under the Receiver Operating Characteristic Curve but instead to search for combinations of parameter values which yield the highest Pearson correlation value *r*^2^ for ST measured over manually digitized landmarks versus ST measured with automatically digitized landmarks.

### Statistical analyses

All statistical analyses were performed using R version 3.4.3 (R Team 2016). We performed two different sets of statistical analyses to investigate how ST changes across species, year of collection and temperature. First, we fitted a standard multiple linear regression with species, year of collection and temperature as numeric predictors using the standard R function lm(). We chose the best model based on the variance explained provided by the *r*^2^ value. Table 2 presents the output of the lm() function using the R package jtools (v1.0.0) (https://cran.r-project.org/web/packages/jtools/jtools.pdf) and its function export_summs() with “scale” and “transform.response” set to “TRUE” which scales and centers the response variable and reports standardized regression coefficients with their heteroskedasticity-robust standard errors. Second, we performed a regression tree analysis and performed cross-validation using recursive partitioning with the regression trees R package “rpart” version 4.1.13 (Therneau *et al.*, 2018) and the associated function rpart() with the “anova” method and obtained the approximate *r*^2^ from a 10-fold cross-validation using the rsq.rpart() function. To confirm the importance of each factor on ST change, we also performed random forest regression analysis using the R package “RandomForest” version 4.6.14 and the randomForest() function in order. Both sets of statistical analyses investigate the role of predictors in explaining a significant part of the variance, multiple linear regression allows the use of interaction terms while regression trees are easier to interpret (James *et al.*, 2013). In addition to these analyses, we systematically plot distribution of ST across predictors (Fig. 2) showing individual values together with mean, standard errors (which directly inform about two-by-two statistical significance between groups), median, quartiles and estimates of the 95% confidence interval of the medians, calculated as 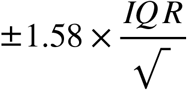 where *IQR* is the interquartile range and *n* the number of individuals for that *IQR* (Chambers *et al.*, 1983).

**Figure 2.**
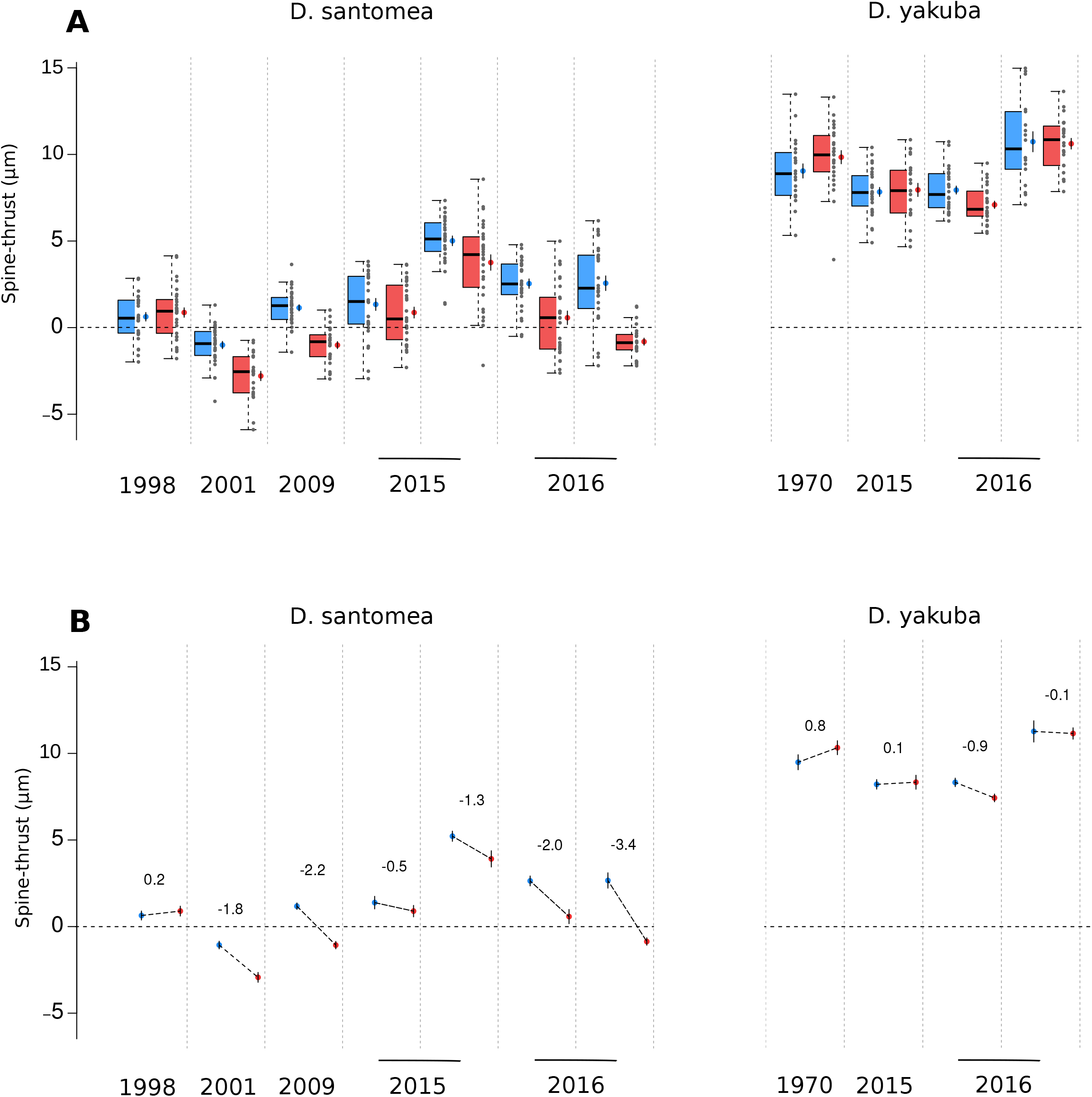
Ventral branches form is sensitive to temperature in *D. santomea*. Isofemale lines arranged by year of collection (see Table 1). (A) For each line, individual values (grey points), median (thick black line), quartiles (colored boxplots), mean (colored points), standard errors (black vertical segment over the colored points) and 95% confidence interval estimates of the median (top and bottom notches) of automatically measured spine thrust are shown. Each line was reared at 18°C (blue) or 25°C (red). (B) For each line, the same mean and standard errors as in panel A are shown, together with the effect slope and corresponding value of that effect (in μm).

**Table 2.** Results of linear model fitting. This best model shows the contribution of each explanatory variable, considered as a numerical value, and their interactions to the overall variance of ST in the full *D. santomea, D. yakuba* dataset (shown in Figures 2A-B) at both temperatures (18°C and 25°C) across all years using the standard R function lm(). The standardized effect values and their heteroskedasticity-robust standard errors are reported together with the range of their *p*-values. For example, species having an overall effect of 1.67 implies that going from *D. santomea* to *D. yakuba* (all other things being equal) increases spine thrust absolute value by a relative (compared to the other effects), dimension-less, value of 1.67. The raw effects together with the full output of the model are provided in Table S1 (see methods). SE: standard error.

| **Factor**<br>*N*=584, *R*<sup>2</sup> = 0.76 | **Effect** | **SE** | ***p*-value** | **Significance** |
| --- | --- | --- | --- | --- |
| Species | 1.67 | 0.06 | < 0.001 | *** |
| Years | 0.54 | 0.06 | < 0.001 | *** |
| Temperature | -0.29 | 0.05 | < 0.001 | *** |
| Species x Years | -0.56 | 0.07 | < 0.001 | *** |
| Species x Temperature | 0.31 | 0.09 | < 0.001 | *** |
| Years x Temperature | -0.23 | 0.09 | < 0.05 | * |

## Results

### Spine thrust (ST) can be measured semi-automatically

We previously reported that the shape of ventral branches in *D. santomea*, *D. yakuba* and their hybrids can be characterized with a set of five manually detected landmarks, which allows to calculate via simple arithmetic how much the lateral spines rise above the central ridge, as a quantitative value named “spine thrust” (ST), expressed in micrometers (Peluffo et al. 2015). The manual positioning of landmarks requires each point to be carefully positioned on the exact feature for the ST measure to be exact. It can introduce between-user and between-sample variability. In particular, the positioning of the three central landmarks can be equivocal and may differ between users.

To use a less biased approach and automate the process, we decided to develop a new measurement method that relies on manually digitized contours of the ventral branches, which are easier to define than landmarks. The position contour of the ventral branches was digitized by hand at an approximately four times faster rate than landmark detection, because it can be done in a single stroke with a digital pen and the resulting ST measure is barely sensitive to the exact pen position. We designed a pipeline that automatically identifies the five landmarks based on the curvature of the manually digitized contours of the ventral branches, and then calculates ST (Fig. 1). The typical rounded form of *D. santomea* is then characterized by a null or negative value of ST (Figure 1, central panel) whereas the spiny form of *D. yakuba* is characterized by a positive value of ST (Fig. 1, right panel). Note that our method does not separate size and shape (Klingenberg, 2016), but considers morphological form as a single quantifiable entity.

To assess repeatability, we digitized twice, at one month interval (at the beginning and at roughly the midpoint of the digitizing effort), 30 individuals of the most characteristic *D. yakuba* strain (Oku, sharp spines) and 31 individuals of the *D. santomea* strain which is the most divergent from this *D. yakuba* strain (1563, extremely rounded shape and small spines). We found a good correspondence between the two sets of automatic measures (Fig. S2), indicating that our pipeline produces robust quantification of ventral branch form.

### Learned *α* and *β*

We find that the same set of 30 *D. yakuba* and 31 *D. santomea* individuals is enough to identify optimal parameter values for *α* (contour smoothing) and *β* (curvature smoothing). We find that with *α* = 0.025 and *β* = 0.055 we obtain *r*^2^ = 0.9 (Figure S1). Although a few other combinations of *α* and *β* yield the same *r*^2^ (Figure S1), we choose this set because it is the one which applies the lowest degree of smoothing.

### Strong interspecific difference in ST

In total, with our semi-automated method (and after removing n = 71 individuals incorrectly dissected or mounted, 12% of total samples, with no apparent distribution bias), we phenotyped 684 individuals raised at 18°C or 25°C throughout their development, corresponding to four *D. yakuba* lines and seven *D. santomea* lines collected between 1998 and 2016 (Table 1). We checked all the automatically detected landmarks by eyes and found that 30 individuals were incorrectly digitized, with a few landmarks either missing or aberrantly positioned (see Fig. S3 for a sample of such individuals), and we excluded these individuals (4% of 684) from subsequent analysis. These aberrant landmark profiles were found in almost all the lines and at both temperatures, with no apparent distribution bias.

At 18°C and 25°C, for *D. santomea* and *D. yakuba*, in all 11 wild isofemale strains, we observed within-strain variability in ST values (Figure 2A, for all groups, *n* per group is between 26 and 31). At both temperatures, the mean ST of each of the seven *D. santomea* strains is inferior to the mean ST of any of the four *D. yakuba* strains (Figure 2A). All *D. yakuba* individuals have a positive ST, while most *D. santomea* strains have a mean ST close to 0 (Fig. 2A). Accordingly, multiple linear regression analysis where the best fit model is ST ~ species x years x temperature shows that the *species* independent variable explains a significant part of the variance in ST (*p*< 0.001, Table 2). Overall, and despite within-strain variability as well as sensitivity to temperature variation, we confirm a morphological difference of ventral branches between wild strains of *D. santomea* and *D. yakuba* using our semi-automatic method of form quantification based on ST (Figure 2A, Table 2).

### Ventral branches of *D. santomea* are plastic to temperature whereas D. yakuba ventral branches are not plastic

For *D. santomea*, in all strains but the oldest one collected in 1998, the mean ST is systematically smaller at 25°C compared to 18°C and standard errors do not overlap (Fig. 2). In contrast, no significant difference in mean ST between 25°C and 18°C is observed for *D. yakuba* strains, except for one strain collected in 2016 (*D. yakuba* Raphia) (Figure 2B). Multiple linear regression analysis supports a negative effect of temperature, as seen with *D. santomea* (p< 0.001, Table 2) and that effect is dependent on species (p< 0.001, Table 2). For the most recently collected wild strain of *D. santomea* (BM16.2), we compared the contours of the two most representative individuals raised at 18°C and 25°C, i.e., the two individuals with ST values closest to the median value of their group. We observed that the individual raised at 18°C has a more *D. yakuba*-like shape of ventral branches compared to the individual raised at 25°C (Fig. 3).

**Figure 3.**
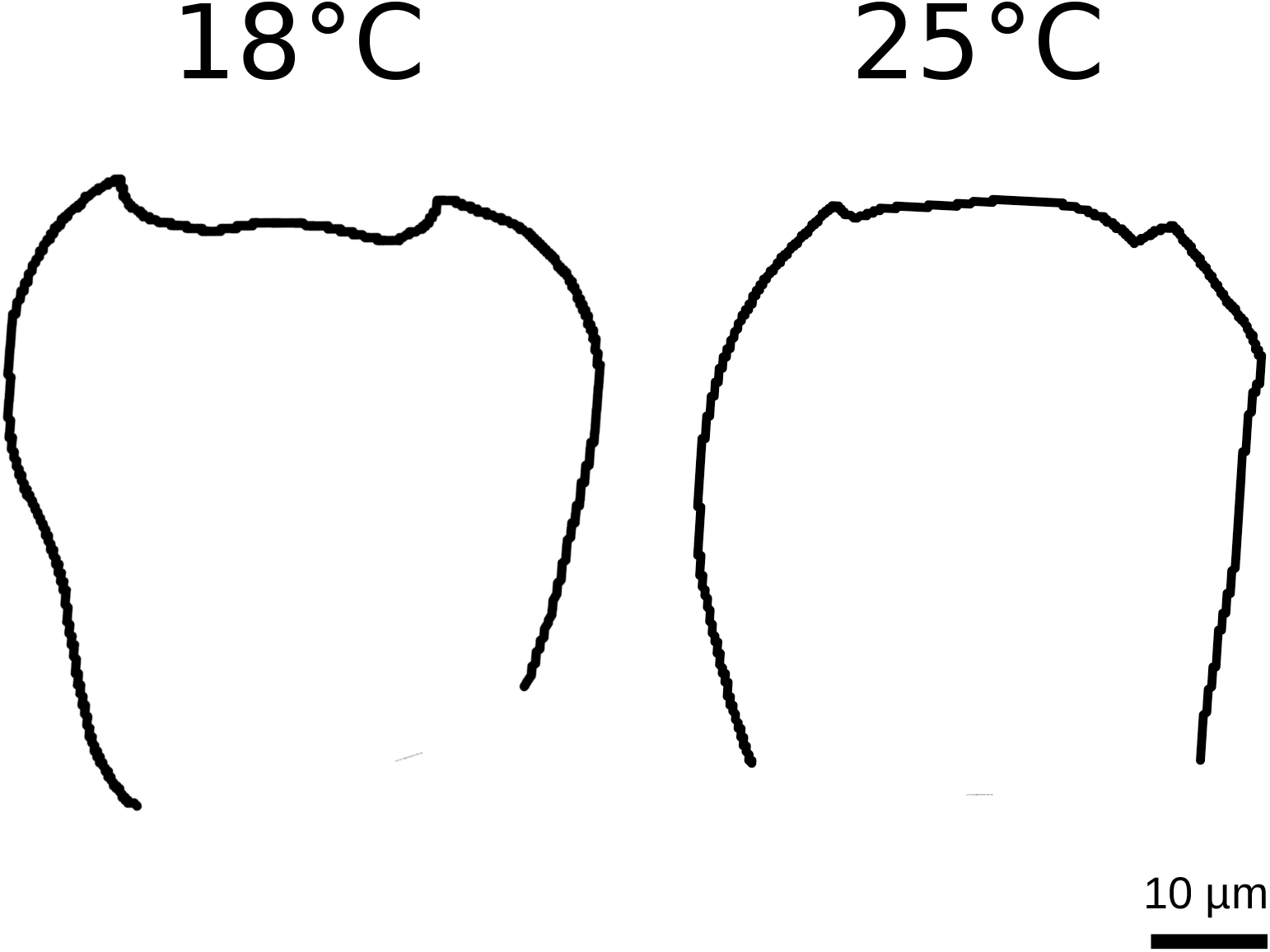
Difference in contour shape at 18°C versus 25°C within the same *D. santomea* isofemale strain collected in 2016. Contours of the two individuals that have the closest spine thrust value to the median value for D. santomea BM16.2 (right most strain on Figure 2A) at 18°C and 25°C.

We find that the statistically significant effect of temperature on *D. santomea* is also statistically dependent on the year at which the strain was collected (p < 0.05, Table 2). In order to interpret our statistical analysis with multiple regression, we performed a 10-fold cross-validated regression tree analysis on the full data set (2 species, 11 strains, 584 individuals). The 10-fold cross-validated error rate is 0.3% and using an additive model of the shape ST ~ species + years + temperature. We found that the variance in the data set is first best partitioned by species and that temperature partitions the data set best for strains collected in 2015 and 2016 (Fig. 4, total variance explained as assessed by cross-validation *r*^2^ is 0.77). To confirm those results, we also performed a random forest regression analysis with the same model as for the regression tree and found that the overall variance explained is *r*^2^ = 0.74 and that the rank of importance of each independent variable is species > years > temperature. Altogether, our results show that in *D. santomea*, but not in *D. yakuba*, ventral branches are sensitive to temperature during development and that this effect is stronger in recently collected strains.

**Figure 4.**
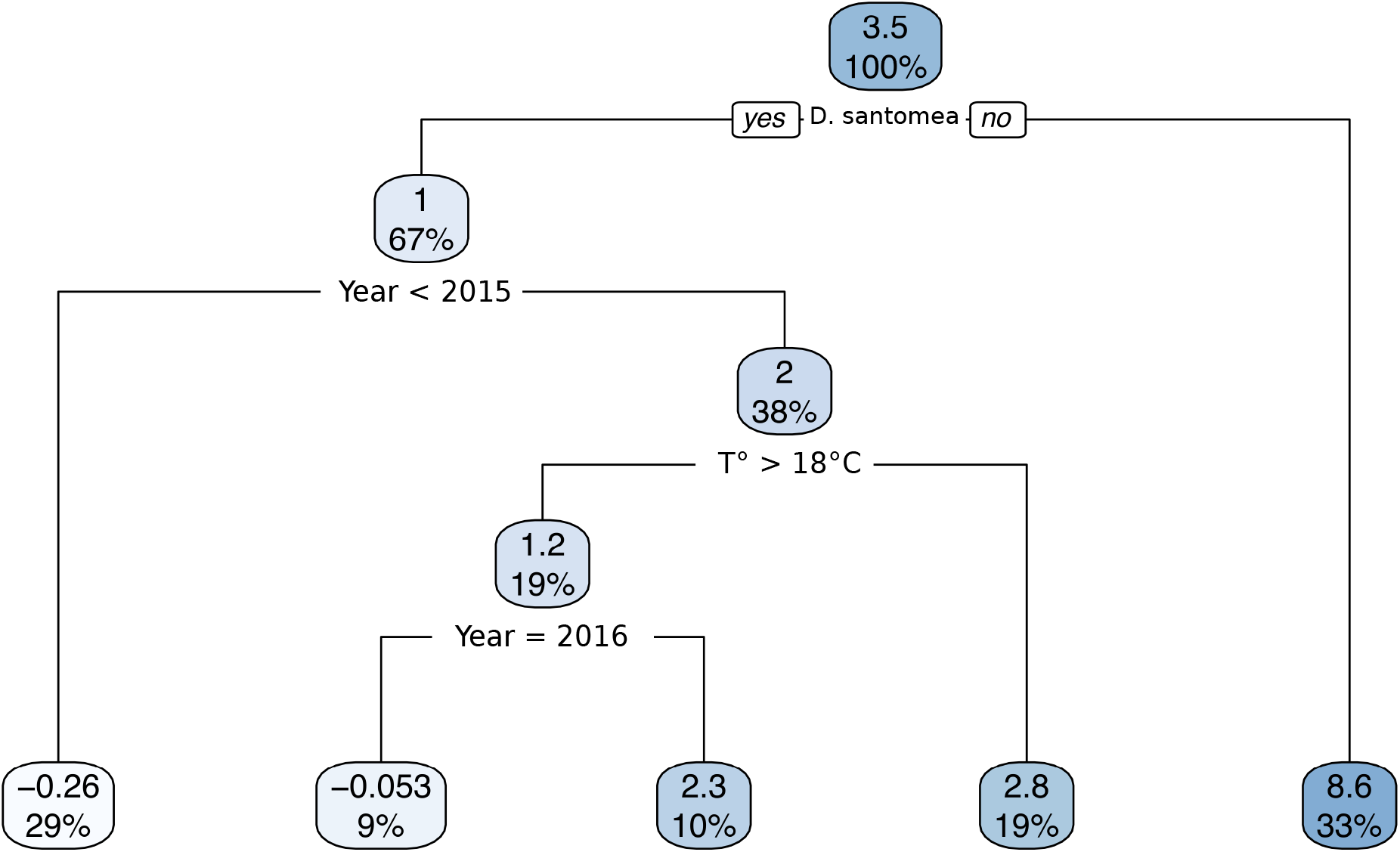
Regression tree for spine thrust measures of all *D. santomea* and *D. yakuba* isofemale strains at both 18 and 25°C. Each node gives the spine thrust mean of all samples included in that node and the proportion of the total dataset included in that node. Below each node are two alternatives: to the left the condition is true and to the right the condition is false. Note that the split between *D. santomea* and *D. yakuba* happens at the top, thereby suggesting that neither temperature nor years have an effect on spine thrust within *D. yakuba*.

### The effect of temperature on Spine Thrust is as high as the effect of the major QTL between *D. yakuba* and *D. santomea*

To compare the effects of temperature and of interspecific genetic variation on ventral branch form, we used our previous QTL mapping dataset of ventral branch form between *D. santomea* and *D. yakuba*, which comprises 365 *D. santomea* backcross individuals (Peluffo *et al.*, 2015). In this previous study, all flies were reared at 25°C as we found that this temperature was optimal to rear both species. The five landmarks were placed manually on images of the ventral branches. A generalized Procrustes analysis was performed on a set of 365 backcross progeny individuals and a larger dataset including the backcross progeny, F1 hybrids and parents. We found that, in both cases, the principal component PC1 explains an important part of the variance (58% in the full dataset and 41% in the backcross), that they are highly correlated (r^2^ = 0.996) and that PC1 in the backcross is highly correlated to ST (spine thrust) (r^2^ = 0.841) and not to centroid size (r^2^ = 0.038).

This QTL mapping study revealed that a 2.7Mb locus on chromosome 3L explains 30% of the mean species difference in ST, meaning that replacing one *D. santomea* allele at this locus with a *D. yakuba* allele leads to an increase in ST of about 3 *μ*m (Peluffo *et al.*, 2015). Pooling all the *D. santomea* lines examined in the present study, we find that a change in the raising temperature from 18°C to 25°C leads to an increase in ST of about 3.4 *μ*m (Fig. 5). We conclude that the effect of temperature is as high as the effect of genetic variation at the major interspecific genetic locus.

**Figure 5.**
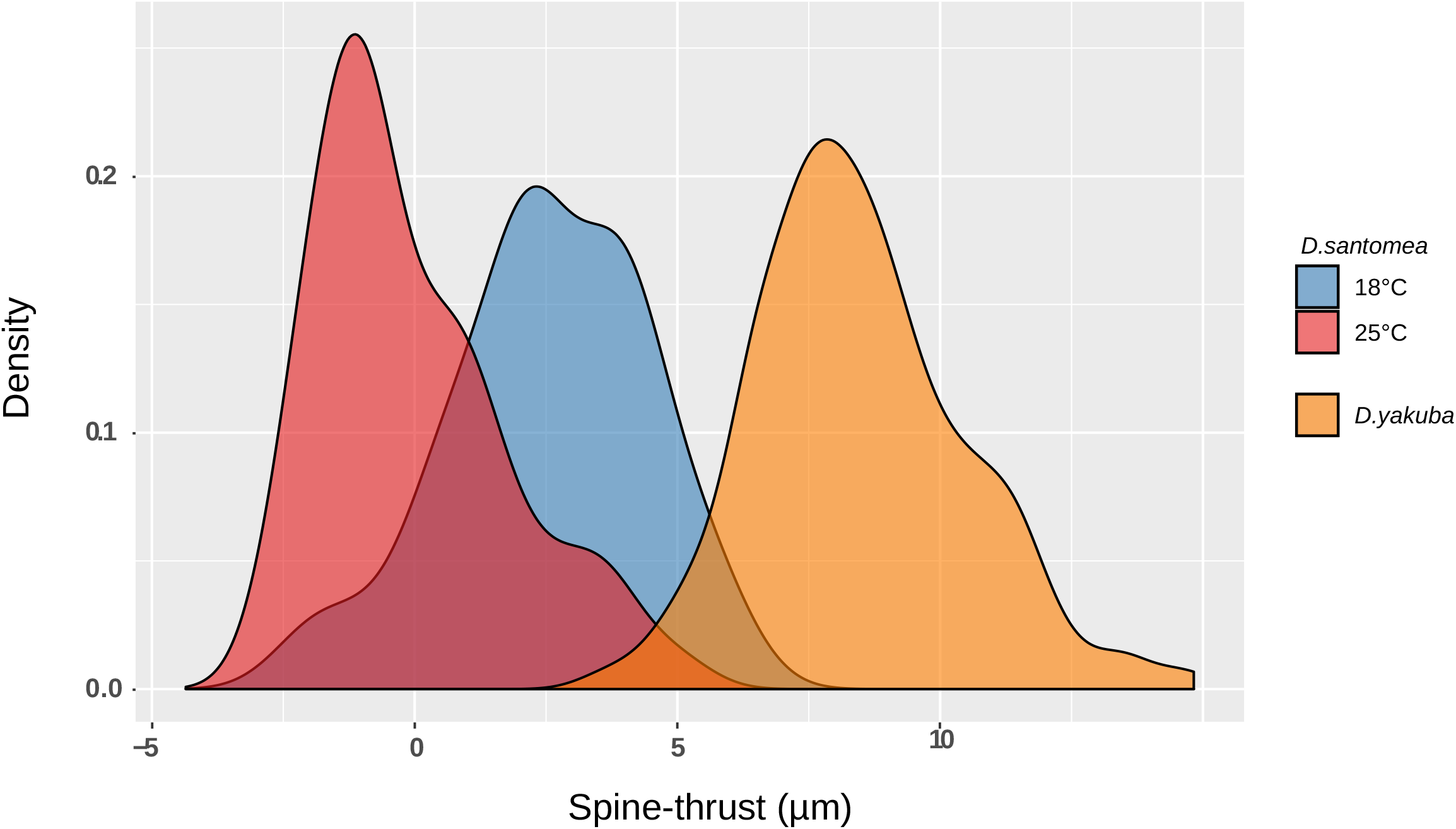
Density distribution of spine thrust values for *D. santomea* lines at 18 and 25°C and *D. yakuba* lines. Density distribution inferred from *D. santomea* males raised at 18°C (blue) and 25°C (red), *D. yakuba* males at both temperatures (orange). Distributions are inferred from the total data shown in Figure 2A.

While the ST density distribution of all *D. yakuba* individuals shows little overlap with the ST density distribution of all *D. santomea* individuals reared at 25°C, it overlaps more with *D. santomea* individuals reared at 18°C (Fig. 5). Overall, our results show that decreasing the temperature from 25°C to 18°C yield *D. santomea* males with spinier, *D. yakuba*-like, ventral branches in the same way as introgressing a *D. yakuba* alleles in place of a *D. santomea* allele at the major interspecific locus.

## Discussion

### A dataset-independent simultaneous quantification of shape and size

Our semi-automatic method, which relies on two simple layers of contour transformation adjusted by regression based learning, is fast and allows the measure of form variation through the simple outlining of ventral branches on 2D pictures. We note that in the future, progress in edge detection algorithms (which for now introduce too much error to measure with precision variations of the order of a few micrometers) might allow full automation from pictures to form quantification.

Having drawn contours, we could also have relied on Fourier based analyses. However, such methods require closed contours which in our case are difficult to draw since the base of the ventral branches is a complex structure which cannot be easily delimited from the cuticle of the ventral branches (Fig. 1). In addition, our method is more suitable for contours in which very large and very small curvatures coexist. Furthermore, an important limitation of morphometrics analyses on landmark data (e.g. Procrustes principal component analysis) is that the PC values are dimensionless (Klingenberg, 2010) and may be difficult to relate to physical features. With our simple measure of ST obtained from the automatically detected landmarks, we are able to quantify and compare forms across studies. Importantly, because we deal with absolute geometric measurements, our method simultaneously analyzes shape and size, unlike most morphometric approaches (Claude, 2008; Klingenberg, 2010). We believe this to be a strength in our case since both shape and size of ventral branches probably contribute to the lock-and-key mechanism; e.g. spiny but short *D. yakuba* ventral branches may not harm *D. santomea* females (Kamimura, 2012; Kamimura & Mitsumoto, 2012b).

### Effect of temperature on size and shape

In most insects and other ectotherms, adult body size typically increases with lower temperatures (Angilletta Jr *et al.*, 2004). Bergmann’s rule, which posits an increasing body size with higher altitude, has been observed within the São Tomé island for the terrestrial caecilian *Schistometopum thomense*, over a temperature range of 9°C (Measey & Van Dongen, 2006). In contrast to other body parts, the genitalia of insects, and of *D. melanogaster* in particular, have been reported as not, or little, plastic in response to temperature or other types of environmental variation (Wheeler *et al.*, 1993; Eberhard, 2009; Shingleton *et al.*, 2009). We find here that this is true also for *D. yakuba* but not for *D. santomea*: changing the rearing temperature from 25°C to 18°C leads to an increase in spine thrust in *D. santomea* male genitalia that is similar to what isobserved between *D. santomea* and *D. yakuba*. In the present study, we only analyzed the effect of temperature on spine thrust and not on the overall shape of the ventral branches. It would be interesting to include additional landmarks to capture the entire shape of the genital structure and to test whether temperature affects the shape of ventral branches in the same way as the interspecific change (Noble et al., 2019).

For each species, we find that strains raised in the same conditions display different averages in ST, showing that the ventral branches form is influenced by genetic factors and is able to evolve.

Plasticity of ventral branches form was detected for all the tested *D. santomea* strains except the one that was maintained for the longest time in the laboratory. Furthermore, the strains collected recently (in 2009, 2015 and 2016) display more pointed ventral branches at 18°C than the ones collected earlier. This suggests that as flies adapt to the laboratory environment, the plasticity of ventral branches form towards temperature tends to be lost and ventral branches tend to be more rounded. Recent studies show that Drosophila flies can adapt to a laboratory environment in 20 generations only, which corresponds to about 8 months (Langmüller & Schlötterer, 2019).

Based on our experiments, we cannot fully rule out plasticity in *D. yakuba*. It is possible that their genital morphology would be altered in external conditions outside of the specific ones that we assayed here. In any case, we find that in our experimental conditions the plasticity of genital form with respect to temperature is higher in *D. santomea* than *D. yakuba*.

### Laboratory observations should be complemented by analysis of wild-caught flies

Tests in the laboratory show that *D. santomea* flies appear to be poorly adapted to high temperatures (Matute *et al.*, 2009). The optimal temperature for larval survival is 21°C for *D. santomea* and 24°C for *D. yakuba.* Furthermore, when adult flies initially raised at 24°C are allowed to distribute themselves along a thermal gradient, they show a preference for 23°C for *D. santomea* and between 26°C and 27°C for *D. yakuba* (Matute *et al.*, 2009). These observations are in agreement with *D. santomea* being collected at higher altitudes than *D. yakuba* in Sao Tomé. However, the reasons why the exact preferred temperature values observed in the laboratory are different from the temperature values measured in the geographic areas of the two species are unknown. Fly collections in Sao Tomé have mostly been done on the north slopes of the island and in these areas *D. santomea* flies are found at an altitude of 1150m or above (Lachaise *et al.*, 2000), which corresponds to temperatures around 18°C or below (climate-data.org, www.worldclim.org/bioclim). However, we note that on the southern slopes of the island a few *D. santomea* flies have also been collected at lower altitudes (650m) in the dense mist forest near Rio Queijo (Matute & Coyne, 2010; Nagy *et al.*, 2018). This suggests that *D. santomea* flies can also inhabit warmer regions of the island and that they might be found across the native forest of Sao Tomé, which goes down to sea level on the western slope of the island (Bell & Irian, 2019). Interestingly, this type of coexistence is not unique on the island: two sister species of frogs closely match the distribution of *D. santomea* and *D. yakuba*, respectively, with the endemic species *Hyperolius thomensis* tied to wet forest habitats while its sister species *H. molleri* is in dry, human-disturbed areas, and *H. thomensis* frogs have also been found in the southern forest at 150 m (Bell & Irian, 2019).

It would be interesting to examine the genitalia of wild caught individual males of *D. santomea* to check the form of their ventral branches at various altitudes. One possibility is that at low altitudes in the southern part of the island *D. santomea* flies display rounded ventral branches while in the hybrid zone with *D. yakuba* at 1150 m and at higher altitude, where temperatures are 18°C or below, they have spinier ventral branches. Of note, *D. santomea* flies have always been collected from traps and have never been observed directly in their native environment: it is possible that they live in microenvironments whose temperature is distinct from the one measured by climate stations (Feder *et al.*, 2000; Negoua *et al.*, 2019).

### Evolution of the plasticity of ventral branches form

To understand the relevance of this temperature sensitivity of genital form for the past and present evolution of *D. santomea* and *D. yakuba*, more needs to be learnt about their ecology and the plasticity of the ventral branches form of their closely related species, *D. teissieri*. Ventral branches are only found in the three species of the *D. yakuba* complex, *D. santomea*, *D. yakuba* and *D. teissieri* (Yassin & Orgogozo, 2013). Since ventral branch form plasticity has not been studied in *D. teissieri*, it is unclear whether this plasticity to temperature is an ancestral trait which has been lost in *D. yakuba* or if it is a novel trait which evolved in *D. santomea* only. The species *D. teissieri* is not found in São Tomé but on the mainland and a few islands of the African continent; it can hybridize with *D. yakuba* (Turissini & Matute, 2017; Cooper *et al.*, 2018). In *D. teissieri* males, the spines are very long and no layer of cuticle is present between them (Kamimura & Mitsumoto, 2012a; Yassin & Orgogozo, 2013). In any case, even if the extent of ventral branch form plasticity in *D. teissieri* was known, it would still be difficult to reconstruct ancestral trait states based on only three species.

The female protective pouches, into which the spiny ventral branches of *D. yakuba* males fit during copulation, were observed in *D. yakuba* but not in *D. santomea* females raised at 21°C and 25°C (Kamimura & Mitsumoto, 2012b; Yassin & Orgogozo, 2013). It would be interesting to check whether such pouches form in *D. santomea* females raised at 18°C, coinciding with the emergence of spiny ventral branches in males. Furthermore, whether more pointed ventral branches in *D. santomea* males due to lower temperatures affects copulation, reproduction and female physiology after mating is unknown.

If we assume that the São Tomé island species *D. santomea* arose from a *D. yakuba*-like ancestor living on the African continent, one can hypothesize that regression in ventral branch size and their plasticity evolved recently in the lineage leading to *D. santomea*. Such a scenario is opposite to the most common view that posits that morphological diversification tends to proceed through losses of plasticity, rather than gains of plasticity [“flexible stem hypothesis” (West-Eberhard, 2003; Schneider & Meyer, 2017), “plasticity-first” model (Levis & Pfennig, 2016)]. It is possible that the decrease in spine thrust that occurred during evolution in the lineage leading to *D. santomea* was accompanied by a gain of ventral branches form plasticity towards temperature. It is unclear whether the plasticity of ventral branches form to temperature is adaptive. More knowledge about the ecology of *D. santomea* and its sister species will be required to elaborate a convincing scenario to interpret the role of the ventral branch form plasticity that we discovered.

## Conclusion

Our data show that genitalia can be plastic to temperature and that this plasticity can evolve coincidentally with speciation. Whereas the sensitivity of insect genitalia shape to temperature or nutrition has been used previously as a proof against the lock-and-key hypothesis (Arnqvist & Thornhill, 1998; Andrade *et al.*, 2005), our work suggests that genitalia can be plastic without rejecting the lock-and-key hypothesis if the environmentally induced changes do not hamper reproduction within each sister species lineage.

## Abbreviations

SE: Standard Error
QTL: Quantitative Trait Locus
ST: spine thrust

## Supplementary Figures

**Figure S1.**
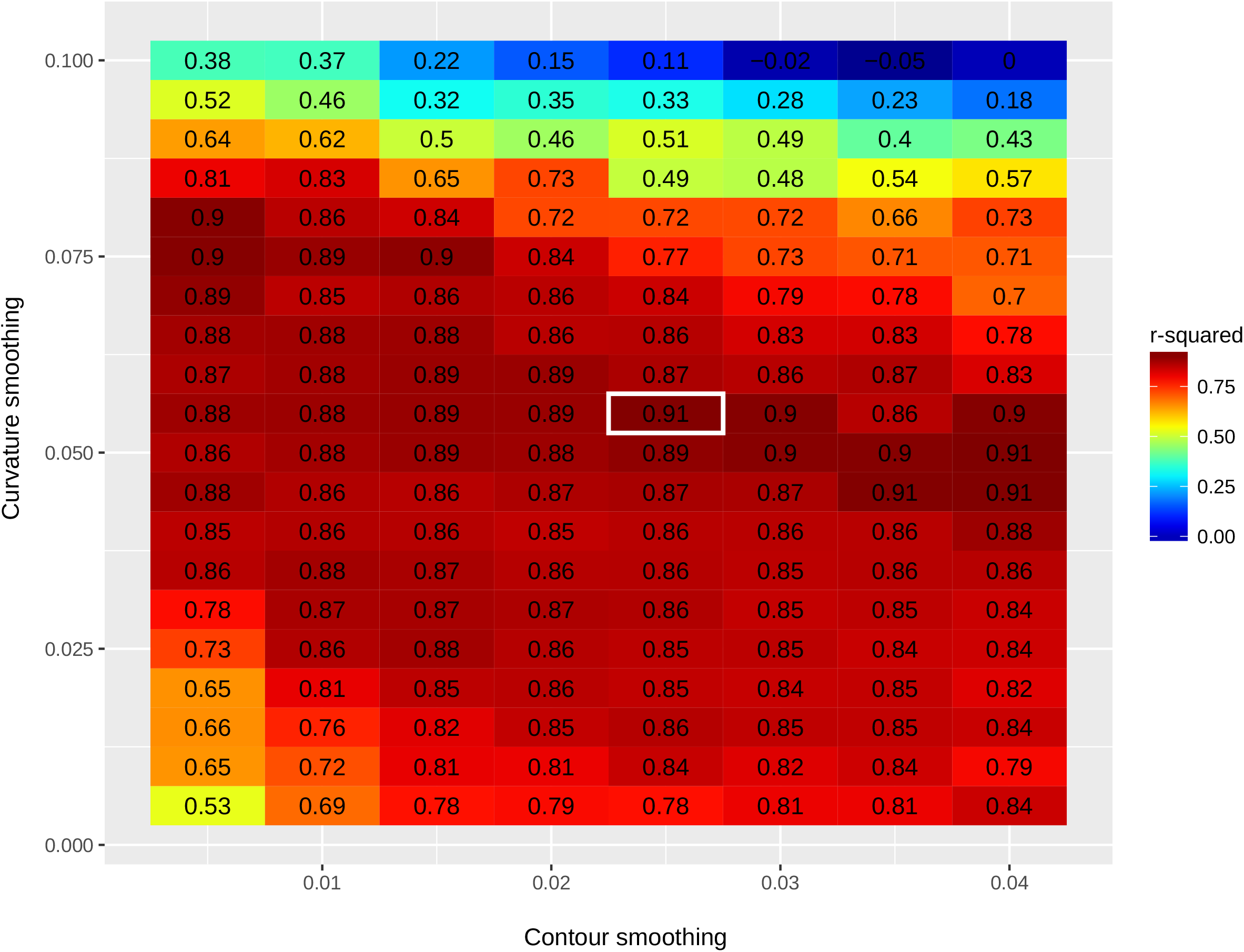
Parameter adjustment for the machine detection algorithm. Training the algorithm relies on two layers of transformation that are each dependent on one parameter: contour coordinate smoothing (horizontal axis) and curvature profile smoothing (vertical axis). Training was performed using a set of 61 individuals, 31 *D. santomea* 1563 and 30 *D. yakuba* Oku (see Table 1) for which we manually digitized both landmarks and contours. For each value of the two smoothing parameters, we performed linear regression of spine thrust from manually digitized landmarks against spine thrust derived from automatically digitized landmarks. The colors and values represent the *r*^2^ from that regression. The value used for all detections is contoured in white.

**Figure S2.**
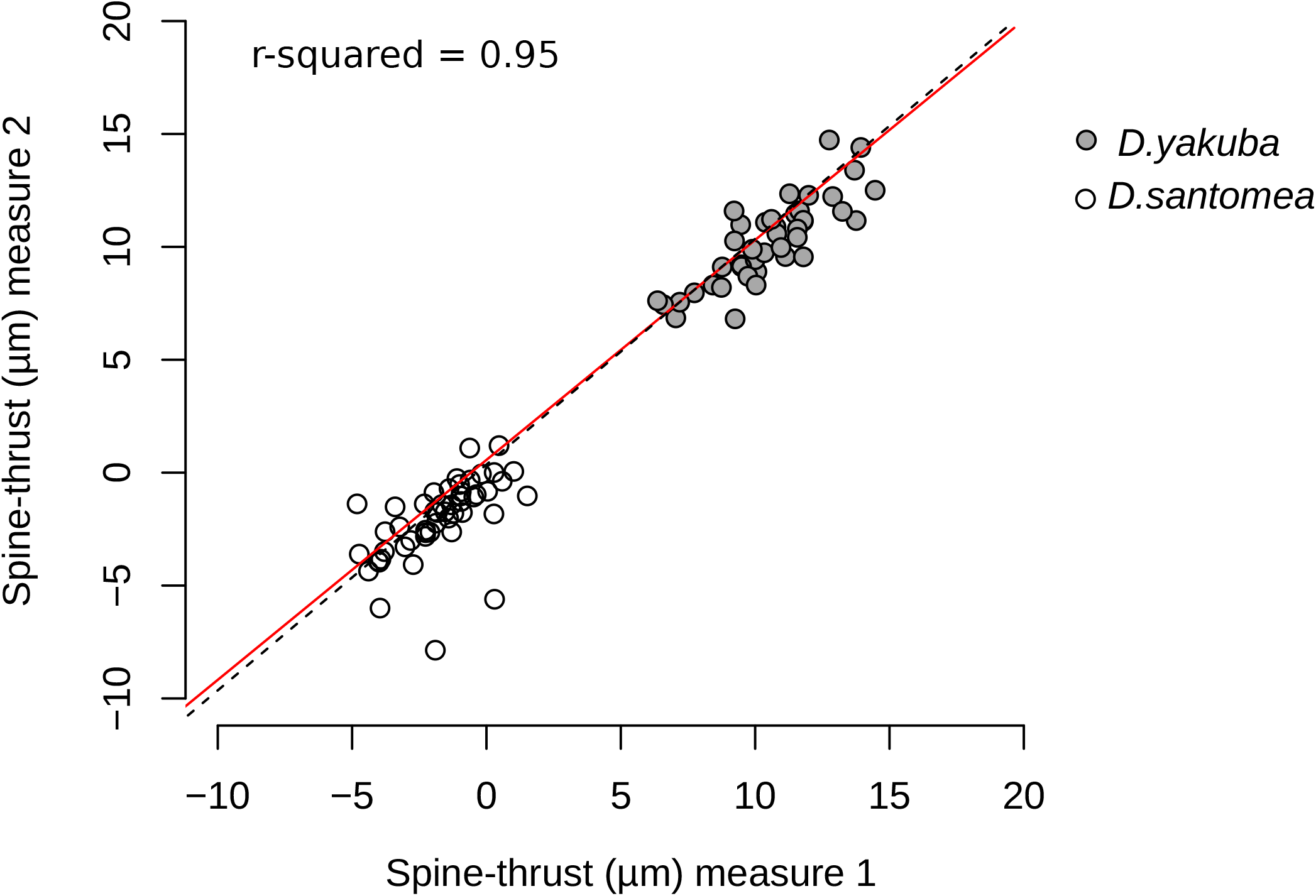
Correlation between two sets of automatic measures from the same dataset. For the training dataset (31 *D. santomea* 1563 and 30 *D. yakuba* Oku), the same user digitized the same contours twice at one month interval and spine thrust was automatically measured. Each point represents one individual. The *y = x* (black dashed line) and linear regression (full red line) are shown.

**Figure S3.**
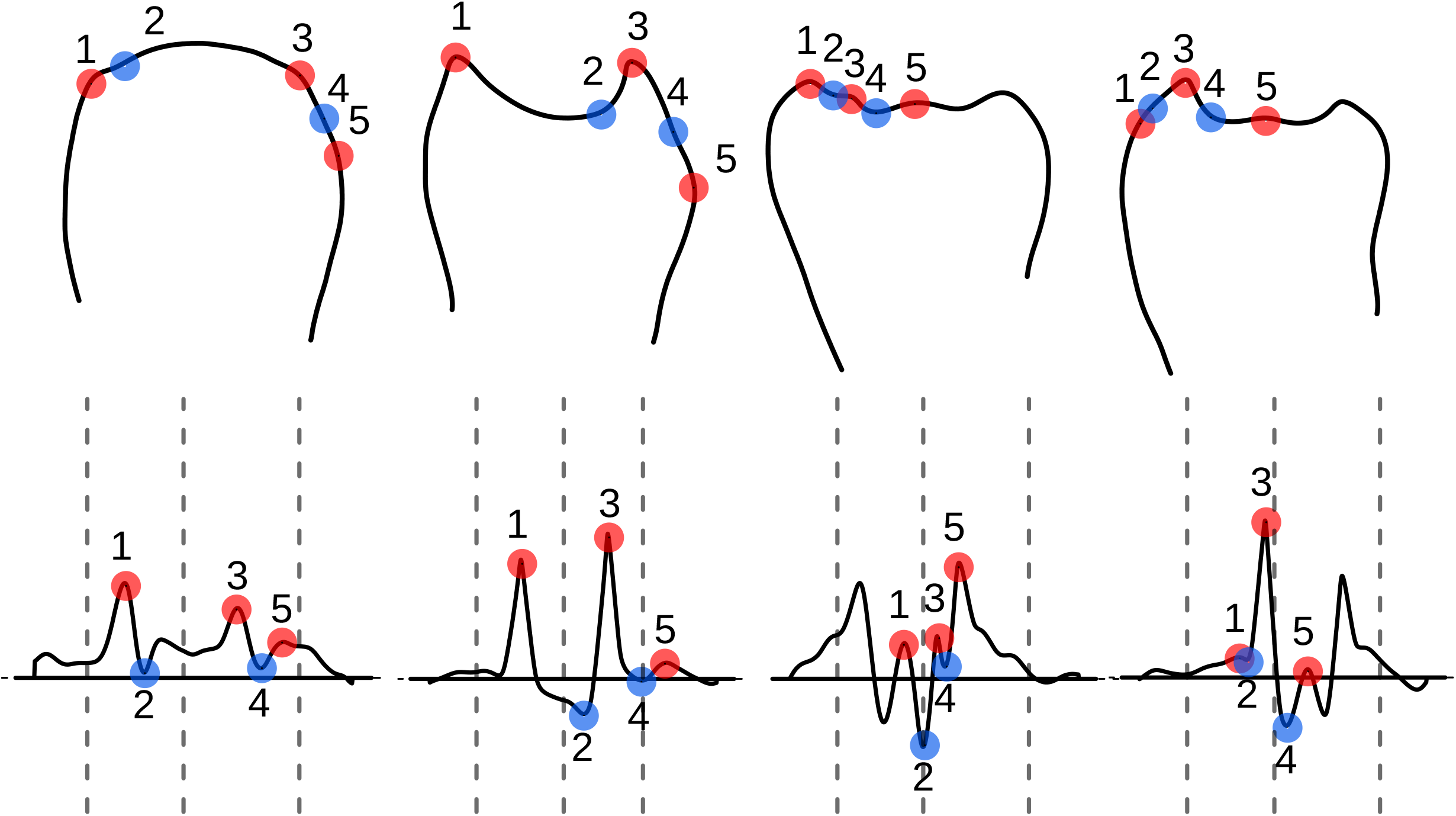
Representative samples of landmarks incorrectly identified with the machine detection algorithm. For each example, we show the smoothed contour, the corresponding curvature profile and identified landmarks.

Table S1. Raw output of R lm (formula = ST ~ Species * Year * Temperature).

## Author Contributions

AEP designed the study, JRD collected fly strains, MH, AVV and AEP collected the data,

AEP and FG designed the analysis pipeline, FM wrote the ImageJ plugin to extract contour pixels, AEP, FG and VCO analyzed the data, AEP, VCO and FG wrote the paper. All authors have read and approved the manuscript.

## Acknowledgments

We are grateful to São Tomé authorities for allowing us to collect flies. We thank David Stern and Daniel Matute for fly strains. We thank the Courtier lab for helpful discussions. We also thank Philippe Rinaudo for feedback on the statistical analyses. The research leading to this paper has received funding from the European Research Council under the European Community’s Seventh Framework Program (FP7/2007–2013 Grant Agreement no. 337579) to VCO and from the labex ‘‘Who am I?’’ (ANR-11-LABX-0071) and the Université de Paris IdEx (ANR-18-IDEX-0001) funded by the French government through grant no. ANR-11-IDEX-0005-02 to AEP.

## DRYAD

doi:10.5061/dryad.kprr4xh1f. The temporary link until manuscript acceptance is https://datadryad.org/stash/share/72YV3b0V2AELCpWuS_ULqMlZyORs9WILt6iMiInAkpE.

---

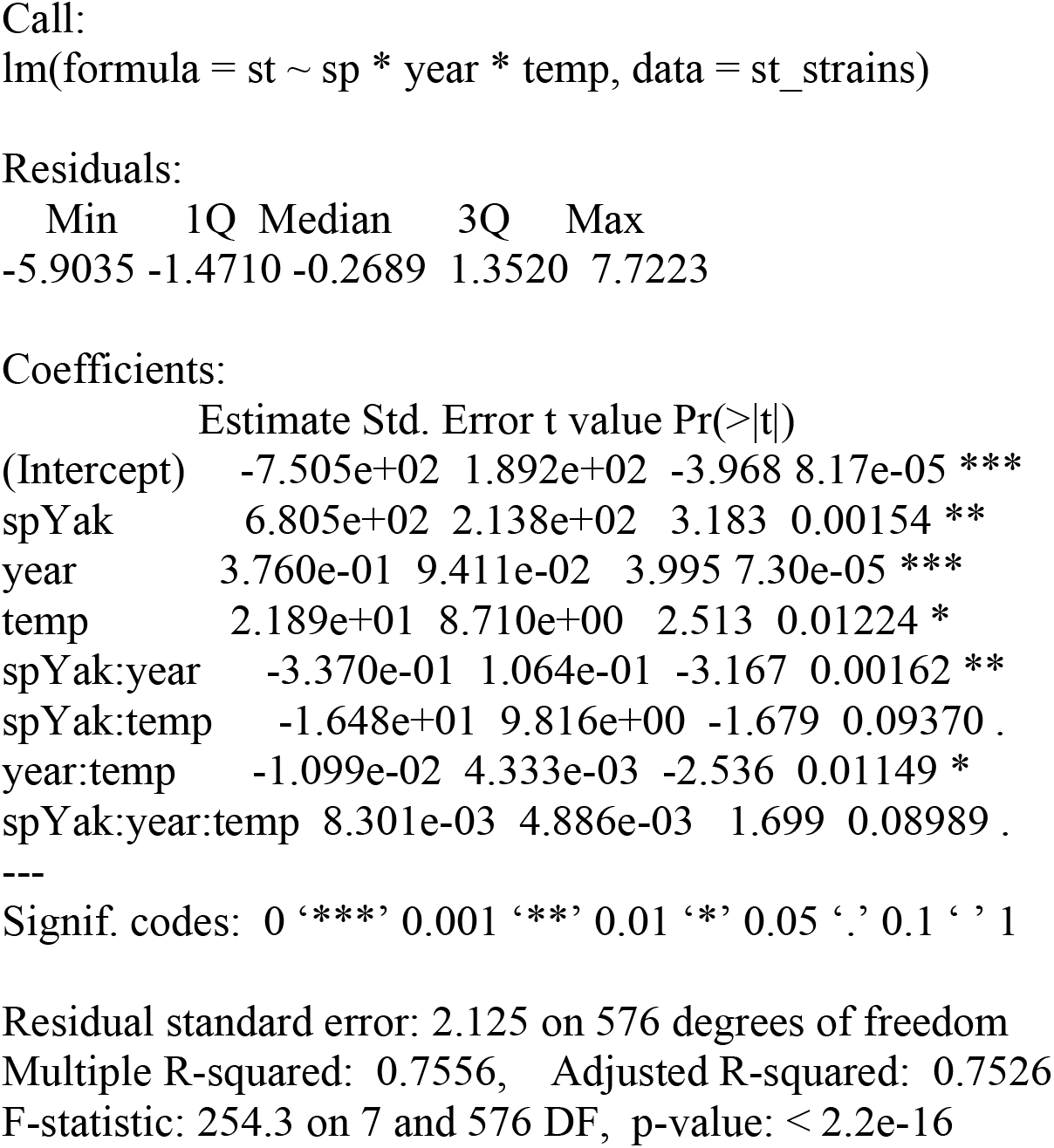

